# FOXM1 Inhibition Promotes Polyploidization and Metabolic Maturation in Human iPSC-Derived Hepatocytes by Modulating the Wnt/β-Catenin Pathway

**DOI:** 10.1101/2025.06.17.660221

**Authors:** Kayque Alves Telles-Silva, Lara Pacheco, Sabrina Komatsu, Fernanda Chianca, Gustavo Chagas, Gabrielly Cristine Martins, Maria Gridina, Daria Panchenko, Valdemir Melechco Carvalho, Elia G. Caldini, Veniamin S. Fishman, Michelle Arkin, Ernesto Goulart, Mayana Zatz

## Abstract

Human induced pluripotent stem cell (iPSCs)-derived hepatocytes are widely used in regenerative medicine and disease modeling. However, existing protocols mainly produce fetal-like cells, limiting accurate modeling of liver functionality. Topoisomerase II (TOP2) and its transcription factor, forkhead box M1 (FOXM1), are silenced during late liver embryonic development; however, their roles in hepatocyte differentiation remain unclear. Here, we examined the effects of TOP2 and FOXM1 inhibition on the terminal differentiation of hepatocytes. We found that subtoxic TOP2 inhibition reduced nuclear chromatin condensation without causing DNA damage. RNA-seq analysis showed that TOP2 inhibition induced cell cycle arrest in a TOP2A-selective manner, with FOXM1 downregulation. ATAC-seq validation demonstrated that TOP2A inhibition decreases chromatin accessibility and modulates the Wnt/β-catenin pathway. Proteomic analysis revealed that FOXM1 inhibition modulated TOP2A expression, replicated TOP2A-mediated cell cycle arrest, and reduced the levels of fetal hepatocyte proteins (HBG1/2, UGT2B7, and AFP). Prolonged FOXM1 inhibition is correlated with increased hepatocyte polyploidization, enhanced CYP450 activity, and improved lipid metabolism, suggesting a potential role in these processes. Overall, our findings suggest that FOXM1 inhibition significantly promotes the terminal differentiation of human iPSC-derived hepatocytes, indicating a potential role for FOXM1 and TOP2A in liver development, regeneration, and disease.

## INTRODUCTION

Human induced pluripotent stem cells (iPSCs) have emerged as a tool to bridge developmental biology, regenerative medicine, and disease modeling because of their capacity to differentiate into various cell types, including hepatocytes (1). The need for a viable hepatocyte source for drug testing has driven the development of new hepatic differentiation techniques (2). Hepatocyte differentiation from iPSCs requires specific factors (Wnt3a, Activin A, HGF) and signaling pathways that control cell proliferation and maturation (3). iPSC-derived hepatocytes show a fetal-like phenotype; thus, they are frequently named hepatocyte-like cells (HLCs), with a proliferative profile and a lack of mature metabolism compared to primary human hepatocytes (PHHs) (4). Therefore, no protocol has achieved terminal differentiation of iPSCs into hepatocytes using either tridimensional, co-culture-based, organoid, or microfluidic platforms (5).

Topoisomerases resolve DNA structures and TOP2 is crucial for DNA replication during embryogenesis (6). TOP2 regulates hepatocyte proliferation (7). Among the TOP2A and TOP2B isoforms, TOP2A upregulation facilitates oncogenesis by enhancing cellular proliferation (8). TOP2A downregulation after weaning coincides with hepatocyte terminal maturation when the liver metabolism changes (9), revealing TOP2A’s role in liver development (10). TOP2A and FOXM1 are frequently associated with hepatocyte proliferation studies (11, 12). FOXM1 regulates cell cycle progression and TOP2A expression during embryogenesis and cancer development (13). The TOP2-FOXM1 interaction maintains a balance between stemness and lineage commitment (14). During late embryonic development, liver parenchymal cells stop proliferating and significantly increase their functionality, especially hepatocyte polyploidization, which is crucial for liver homeostasis (15). Foxm1-deficient mice exhibit reduced hepatocyte mitosis and polyploidy (16), suggesting that FOXM1 may be involved in the regulation of hepatocyte cell cycle and polyploidization. However, the TOP2-FOXM1 axis has not been explored in the context of hepatocyte terminal differentiation.

In this study, we investigated the effects of TOP2 and FOXM1 on human iPSC-derived hepatocyte differentiation. FOXM1 inhibition promoted hepatocyte maturation, including cell cycle arrest and polyploidy. FOXM1 suppression reduced fetal markers while increasing adult hepatic gene expression by modulating the Wnt/β-catenin pathway. TOP2A functions as an integral partner in this network.

## METHODS

### Ethics approval

All the experiments using human iPSCs were approved by Ethics Committees of Biosciences Institute of University of Sao Paulo (USP) (Protocol 1.294.118).

### Treatment with small molecules

For topoisomerase II inhibition, 10 mM etoposide (Sigma-Aldrich) was dissolved in dimethyl sulfoxide (DMSO). For FOXM1 inhibition, 2 mM RCM-1 (Sigma-Aldrich) was dissolved in dimethyl sulfoxide (DMSO) and Etoposide or RCM-1 was diluted in DMSO at 100x a final intended concentration. Next, 100x etoposide or RCM-1 was directly added to the hepatocyte maturation medium and diluted 1:100. Etoposide was predominantly used at a final concentration of 10 μM, while RCM-1 was used at a final concentration of 1 μM. The control groups were treated with 1% DMSO in hepatocyte maturation medium. The medium was changed every 48h, and the treatments were started on differentiation day 20.

### RNA-extraction, RT-qPCR, and RNA-sequencing analysis

RNA was extracted using a PureLink RNA Mini Kit (Thermo Fisher Scientific). Reverse transcription was performed using the Superscript IV Synthesis Kit (Thermo Fisher Scientific). RT-qPCR was performed using Power SYBR Green PCR Master Mix (Thermo Fisher) on a QuantStudio 5 System. The samples were amplified using primers designed using Primer-BLAST or retrieved from PrimerBank (Supplementary Table 1). Expression analysis was performed using the ΔΔCt method and normalized to the RPLP0 or GUSB controls. Human Plateable Hepatocytes were used as the PHH controls and were harvested for 24h. RNA sequencing was performed using the Novogene software. RNA quality was assessed using Bioanalyzer 2100 (Agilent). Library preparation was performed using the TruSeq Stranded mRNA Kit, and sequencing was performed using NovaSeq 6000 (Illumina). The reads were aligned to the human genome hg38 using Hisat2. Differential gene expression analysis was performed using the DESeq2 package in R. The thresholds for differential expression were adjusted P-value ≤ 0.05, and |log2(fold change)| ≥ 1. ClusterProfiler was used to test the enrichment of differentially expressed genes. Data visualization was performed using the clusterProfiler and pheatmap packages and the MaGIC Volcano Plot Tool. PPI analysis was performed using the STRING database and term enrichment was performed using the Enrichr web tool.

### ATAC-sequencing analysis

ATAC sequencing was conducted through Novogene. Nuclei were incubated with a Tn5 transposase mix at 37°C for 30 minutes. Libraries underwent PCR amplification, were purified using AMPure beads, and evaluated with Qubit. Sequencing was performed on the Illumina Hiseq Platform, producing 150 bp paired-end reads. Reads were trimmed with Skewer, aligned using BWA, and filtered for quality (MAPQ 13) and proper pairing (>18 nt). Peak calling was executed with macs2 using the command ‘macs2 callpeak --nomodel --keepdup all-call summits (v2.1.2). Term enrichment was analyzed with Enrichr, and motifs were identified using HOMER. The ATAC-seq library preparation adhered to the ATAC 2.0 protocol. Cells were fixed with paraformaldehyde, quenched with glycine, and stored at −80°C. Following lysis, cells were digested with DpnII, labeled with biotin-14-dCTP, and ligated. Libraries were prepared using the KAPA HyperPlus Kit with biotin enrichment. Data processing was carried out with Juicer and Cooltools. Normalized maps were analyzed using FANC, and compartmentalization was assessed with cooltools saddle functions. A/B compartments were assigned through eigenvector decomposition.

### Proteomic analysis

For proteomic analysis, samples were lysed using buffer containing 50 mM HEPES (pH 8), 1% SDS, 1% sodium deoxycholate, 5 mM EDTA, and 50 mM NaCl, with Halt Protease and Phosphatase Inhibitor Cocktail. Proteins were quantified using the Pierce BCA Protein Assay Kit and 250 μg in 100 μL were digested using a modified semiautomated SP3 protocol. Phosphopeptides were enriched using titanium dioxide beads (Titansphere, Gl Sciences, Japan) according to the manufacturer’s instructions. LC-MS/MS analyses were achieved in a nano LC UltiMate 3000 system connected coupled to an Exploris 240 hybrid quadrupole-orbitrap tandem mass spectrometer (Thermo Fisher Scientific, Bremen, Germany) equipped with an EASY-Spray source, operating in positive ion mode. Spectral data were searched against an in silico human (Uniprot Swissprot database downloaded on August 06, 2024 which contained 20,447 entries) spectral library using DIA-NN 1.9.1 software. FragPipe-Analyst was used for visualization, Enrichr for term enrichment, and the X2K web tool for regulatory networks.

### Flow cytometry analysis

For flow cytometry analysis, cells were dissociated using TrypLE Express Enzyme (Thermo Fisher Scientific). The cells were fixed and permeabilized using the Fix & Perm Cell Fixation and Permeabilization Kit (Thermo Fisher Scientific), according to the manufacturer’s instructions. The samples were blocked with 5% BSA in PBS (1x) for 1 h and then incubated with primary antibodies (listed in Supplementary Table 3) at 4°C for 1h. After washing with PBS (1x), the samples were incubated with Alexa Fluor-conjugated secondary antibodies (Supplementary Table 3) at 4°C for 1 h. samples were analyzed using a FACS Aria III flow cytometer (BD Biosciences) with 20,000 events per group. For ploidy analysis, samples were dissociated with TrypLE Express (Thermo Fisher), fixed with 70% ethanol at -20°C for 2h, and DNA was stained using FxCycle PI/RNase Staining Solution (Thermo Fisher) with 100 U/μL RNase I for 30 min at 37°C, acquiring 500,000 cells per group. For DNA fragmentation and damage assays, we used the Click-iT TUNEL Alexa Fluor 488 kit and anti-phospho-H2AX antibody (Supplementary Table 3). For senescence evaluation, the CellEvent Senescence Green Flow Cytometry Assay Kit (Thermo Fisher Scientific) was used. Data were analyzed using FlowJo V10 software.

### Toxicity assays

For the cytotoxicity assays, HLCs were differentiated into 96-well plates. To assess cell viability, intracellular ATP levels were measured using the CellTiter-Glo Luminescent Cell Viability Assay Kit (Promega, G7571) according to the manufacturer’s instructions. To assess toxicity, 200 μL of the supernatant was collected and lactate dehydrogenase (LDH) was measured using the CyQUANT LDH Cytotoxicity Assay (Thermo Fisher, C20301), according to the manufacturer’s instructions. Both assays were normalized using a blank negative control (hepatocyte maturation medium) and death-positive control (5% Triton-X100 treatment for 15 min).

### Quantification and statistical analysis

Statistical analyses were performed using GraphPad Prism 8 with an unpaired two-tailed Student’s t-test, one-way analysis of variance (ANOVA), and post-hoc Tukey’s test. Statistical significance was set at P < 0.05. n represents the number of biological replicates and independent experiments. G*Power software was used to determine the minimum sample size for data collection to achieve α = 0.05 and power =p0.8.

### Data availability

The RNA-seq, ATAC–seq, and Hi-C data reported in this study have been deposited in the NCBI Gene Expression Omnibus (GEO) with the accession number GSE298619, GSE298820, and GSE298819, respectively. The proteomic data reported in this study have been deposited in the ProteomeXchange Consortium via the PRIDE partner repository under the accession number PXD064121.

## RESULTS

### Subtoxic Etoposide Treatment Promotes Quiescence of HLCs

To assess the effect of etoposide on TOP2 inhibition in HLCs, human iPSCs were differentiated into HLCs (Supplementary Fig. S1A). The HLCs exhibited hepatocyte morphology on day 20 (Supplementary Fig. S1B), and more than 71.4% double-positive staining for ALB and UGT1A1 (Supplementary Fig. S1C) and significant hepatocyte-specific gene expression (*HNF4A*, *ALB*, and *CYP3A4*) (Supplementary Fig. S1D). To determine its subtoxic concentrations, we exposed HLCs to etoposide for five days (Fig. 1A). Etoposide did not induce apoptosis at concentrations up to 3 mM, with similar responses observed in HLCs and HepG2 cells. (Fig. 1B). We also validated apoptosis using teniposide, which is non-toxic at low concentrations (Fig. 1C). Therefore, 10 µM etoposide was selected for further experiments. Etoposide-treated cells showed polygonal morphology similar to that of primary human hepatocytes (PHHs) (Fig. 1D). Similar to the adult PHH organoids, etoposide-treated HLCs lacked strong nucleolar staining (Fig. 1E). At 10 µM, etoposide increased mitochondrial cristae elongation (Fig. 1F), resembling metabolically active PHHs, whereas the control HLCs showed mitochondrial vacuolization. TUNEL and comet assays revealed no significant differences in DNA damage between the treated and control groups (Fig. 1G and H). Flow cytometric analysis of H2AX phosphorylation showed no differences between groups (Fig. 1I). These results confirm that 10 µM etoposide is non-toxic and does not cause DNA damage in HLCs.

**Fig. 1:**
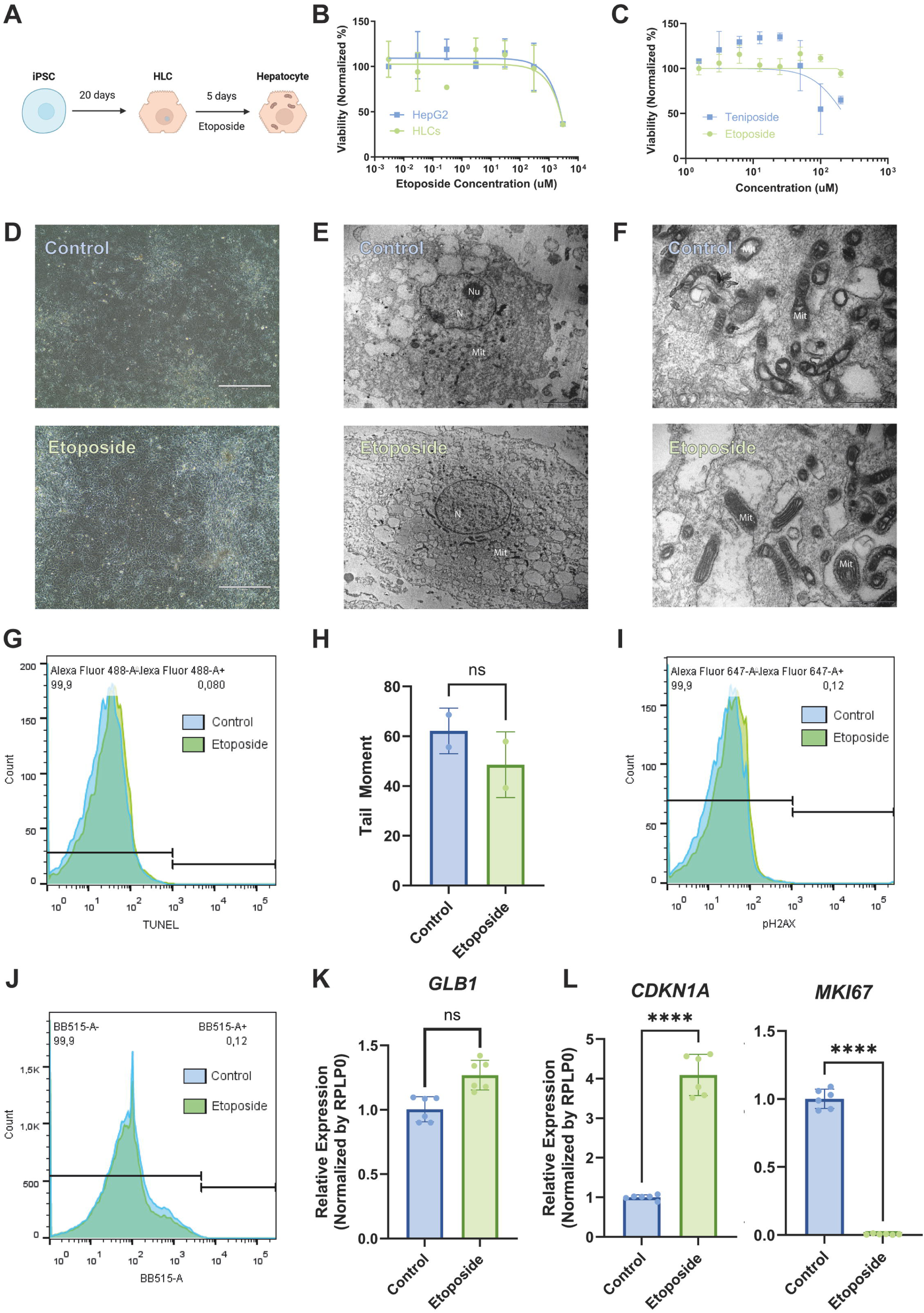
Low-dose etoposide preserves the viability and DNA integrity of HLCs. **A**) Schematic of etoposide treatment of HLCs (Created using BioRender.com). **B**) Cell viability assay of HLCs and HepG2s negative control) with different concentrations of etoposide to determine toxic doses (*n* = 2 biological replicates and 3 independent experiments for HLCs, *n* = 3 independent experiments for HepG2). **C**) Cell viability assay of HLCs with different concentrations of etoposide and teniposide to compare toxic doses. **D**) Brightfield images of HLCs treated with 10 μM etoposide (scale bar = 400 μm). **E**) Transmission electron micrograph of etoposide-treated HLCs (scale bar = 5 μm). N = nucleus; Nu = nucleoli; Mit = mitochondria. **F**) Transmission electron micrograph of etoposide-treated HLCs’s mitochondria (scale bar = 1 μm). Mit = mitochondria. **G**) TUNEL assay of etoposide-treated HLCs. **H**) COMET assay of etoposide-treated HLCs. **I**) Flow cytometry of pH2AX staining of etoposide-treated HLCs. **J**) Flow cytometry of hydrolyzed β-galactosidase staining of etoposide-treated HLCs. **K**) RT-qPCR of *GLB1* of etoposide-treated HLCs. **L**) RT-qPCR of *CDKN1A* and *MKI67* of etoposide-treated HLCs. In every experiment, *n* = 2 biological replicates and 3 independent experiments. In B, C, H, K, and L, data is mean ± SD. In B and C, paired t-test (p<0.05). In H, K, and L, t-test (p<0.001).

We further evaluated the effect of 10 µM etoposide on HLC senescence by measuring the β-galactosidase activity using flow cytometry and RT-qPCR. No significant differences were observed between the treated and control groups (Fig. 1J and K), indicating that etoposide does not induce senescence. Quiescence is relevant in iPSC-derived hepatocytes, marking terminal differentiation in disease modeling and drug development (17). *CDKN1A* expression increases during both cellular senescence and quiescence, whereas *MKI67* expression is downregulated in quiescent hepatocytes. RT-qPCR showed that 10 µM etoposide increased *CDKN1A* and decreased *MKI67* expression (Fig. 1L). Overall, 10 µM etoposide did not induce apoptosis, DNA damage, or senescence in HLCs but potentially induced quiescence.

### RNA-Seq Reveals Etoposide-Induced Transcriptome Rewiring and Cell Cycle Arrest in HLCs

Low-dose etoposide facilitates osteogenesis in certain cell lines (18) and shifts apoptotic pathways from caspase-dependent to differentiation-linked pathways (19). To further evaluate the effect of etoposide on HLCs, RNA sequencing was performed. Intra-group correlations were higher than inter-group correlations, with control and treated HLCs being similar to PHHs (Fig. 2A). PCA showed that etoposide changed the expression profile of HLCs compared to the controls (Fig. 2B), with PC1 accounting for cell source variance (41.25%) and PC2 accounting for etoposide treatment (27.3%). A Venn diagram revealed that 3,301 genes were exclusively expressed in etoposide-treated HLCs (Fig. 2C). Etoposide induced downregulation of global gene expression (Fig. 2D), with *TOP2A* being the most downregulated. Although etoposide inhibited both the TOP2A and TOP2B isoforms, *TOP2B* expression remained unchanged at 10 µM. Among the 1,010 differentially expressed genes, etoposide-induced changes made HLCs resemble PHHs (Fig. 2E). These genes were associated with chromosome segregation, nuclear division, and cell cycle arrest (Fig. 2F). Although etoposide typically causes cell cycle arrest via DNA damage (20), this was not observed in HLCs. Analysis showed that etoposide-treated HLCs resembled PHHs more than control HLCs (Fig. 2G), with downregulation of cell cycle progression genes. Seventeen genes formed the predicted protein interaction core (Fig. 2H). ChEA analysis identified FOXM1 and E2F4 as the main enriched transcription factors (Fig. 2I). FOXM1 regulates cell cycle progression (21), suggesting an involvement of FOXM1 in the maturation of HLCs.

**Fig. 2:**
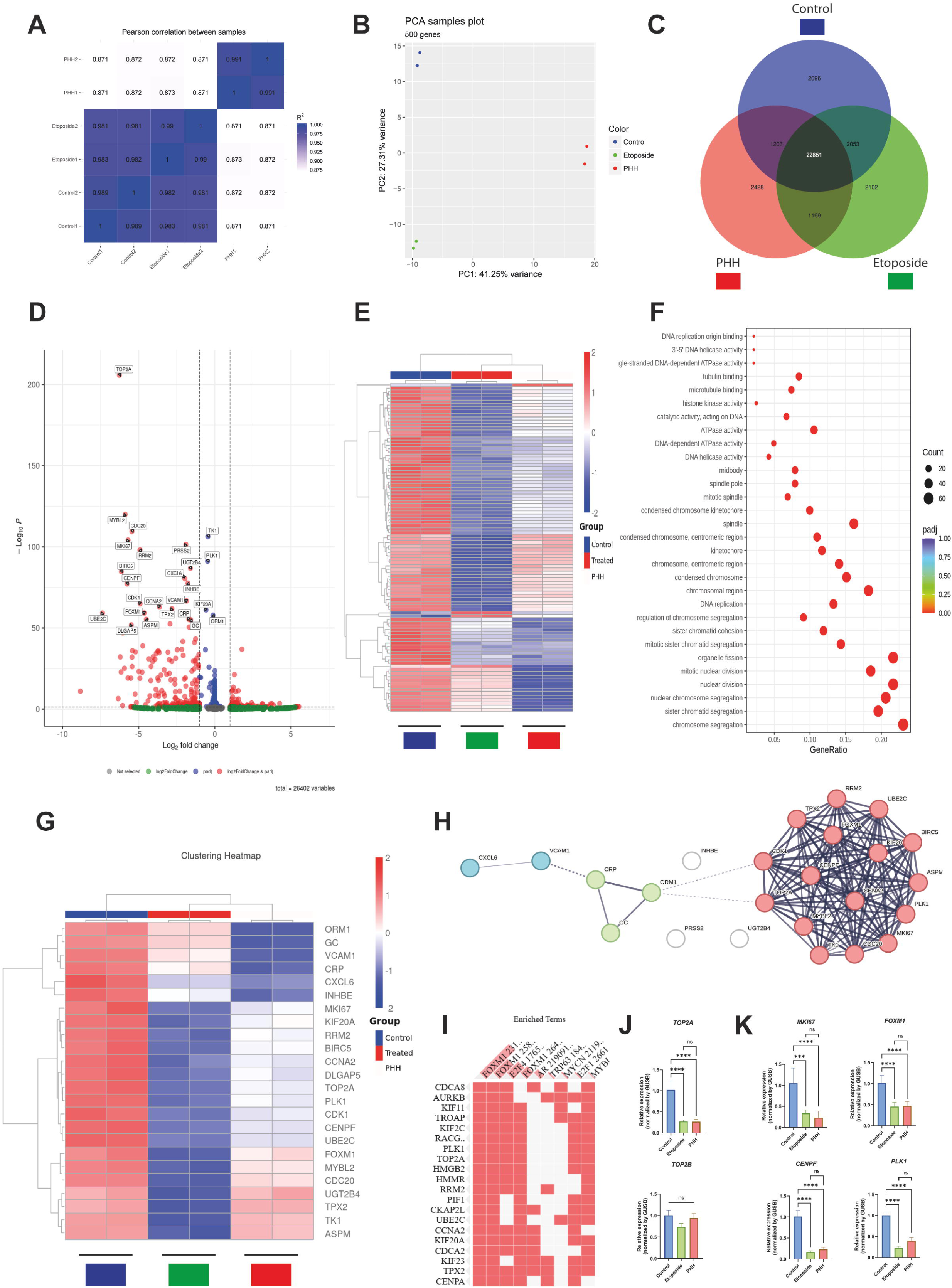
RNA-seq analysis of etoposide-treated HLCs. **A**) Inter-sample correlation matrix heatmap (R^2^ = square of Pearson correlation coefficient) of control HLCs, etoposide-treated HLCs, and primary human hepatocytes (PHHs). **B**) Principal component analysis (PCA) on fragments per kilobase per million mapped fragments (FKPM). **C**) Venn diagram on all identified genes. **D**) Volcano plot of differentially expressed genes between etoposide-treated and control HLCs (|log2(FoldChange)| >= 1 & padj <= 0.05). **E**) Heatmap of the top 100 z-scored differentially expressed genes between etoposide-treated and control HLCs. **F**) Gene ontology (GO) scatter plot for biological processes of the top 100 z-scored differentially expressed genes between etoposide-treated and control HLCs. The size of the circle indicates the percentage of genes in each annotation category. **G**) Heatmap of the top 25 z-scored differentially expressed genes between etoposide-treated and control HLCs. **H**) STRING interactome of the top 25 differentially expressed genes between etoposide-treated and control HLCs. The colours represent k-means clustering. **I**) ChIP-X enrichment analysis (ChEA) of the top 25 differentially expressed genes between etoposide-treated and control HLCs. **J-K**) RT-qPCR of *TOP2A* and *TOP2B* (**J**); or *MKI67*, *FOXM1*, *CENPF*, and *PLK1* (**K**) of etoposide-treated HLCs. In every experiment, *n* = 2 biological replicates. In E and G, the colour represents the average expression level for the indicated gene. In J and K, data is mean ± SD, analyzed by one-way ANOVA with multiple comparisons and Tukey’s correction (p<0.001); *n* = 2 biological replicates and 3 independent experiments.

Next, to characterize cell cycle arrest caused by etoposide in HLCs, we found that 10 µM etoposide showed isoform specificity and reduced *TOP2A*, but not *TOP2B*, gene expression (Fig. 2J). Cell cycle arrest was confirmed by reduced *MKI67* and *FOXM1* expression levels, similar to those observed in PHHs. Downregulation of *CENPF* and *PLK1* expression (Fig. 2K) correlated with hepatocyte polyploidization during embryonic liver development (22), indicating that TOP2A inhibition may induce cytokinesis failure and hepatocyte polyploidization. Overall, RNA-seq indicated that treating HLCs with etoposide shifted their transcriptional profile towards PHHs, especially regarding cell cycle regulation gene sets.

### TOP2A Inhibition Remodels Chromatin Architecture to Suppress Wnt/JAK-STAT Signaling and Cell Cycle Gene Expression

To assess the impact of TOP2A inhibition on chromatin accessibility, we conducted ATAC-seq on etoposide-treated HLCs and their controls. Etoposide is known to induce global chromatin remodeling and transcriptional silencing (23). In etoposide-treated HLCs, we observed an increase in short-range promoter peak identification (Fig. 3A) but a decrease in overall chromatin accessibility (Fig. 3B). The volcano plot illustrated this reduced chromatin accessibility (Fig. 3C), which accounts for the transcriptional repression observed in the RNA-seq analysis (Fig. 2D). Hierarchical clustering revealed a clear distinction between treated and control groups, with treated HLCs exhibiting less accessible peaks (Fig. 3D), aligning with gene silencing (Fig. 2E). KEGG analysis indicated alterations in the Wnt and JAK-STAT pathways (Fig. 3E), which are crucial for fetal hepatocyte proliferation (10). The PI3K-Akt pathway influences the effects of Wnt signaling (24). Post-treatment, chromatin accessibility to PI3K-Akt pathway effectors increased (Fig. 3F). CTCF, linked to Wnt signaling (25), emerged as the most differentially enriched motif between the groups (Fig. 3G). Changes in CTCFL expression suggested reduced binding (Fig. 3H). IGV visualization confirmed decreased accessibility of gene promoters related to hepatocyte proliferation and polyploidization (Fig. 3I).

**Fig. 3:**
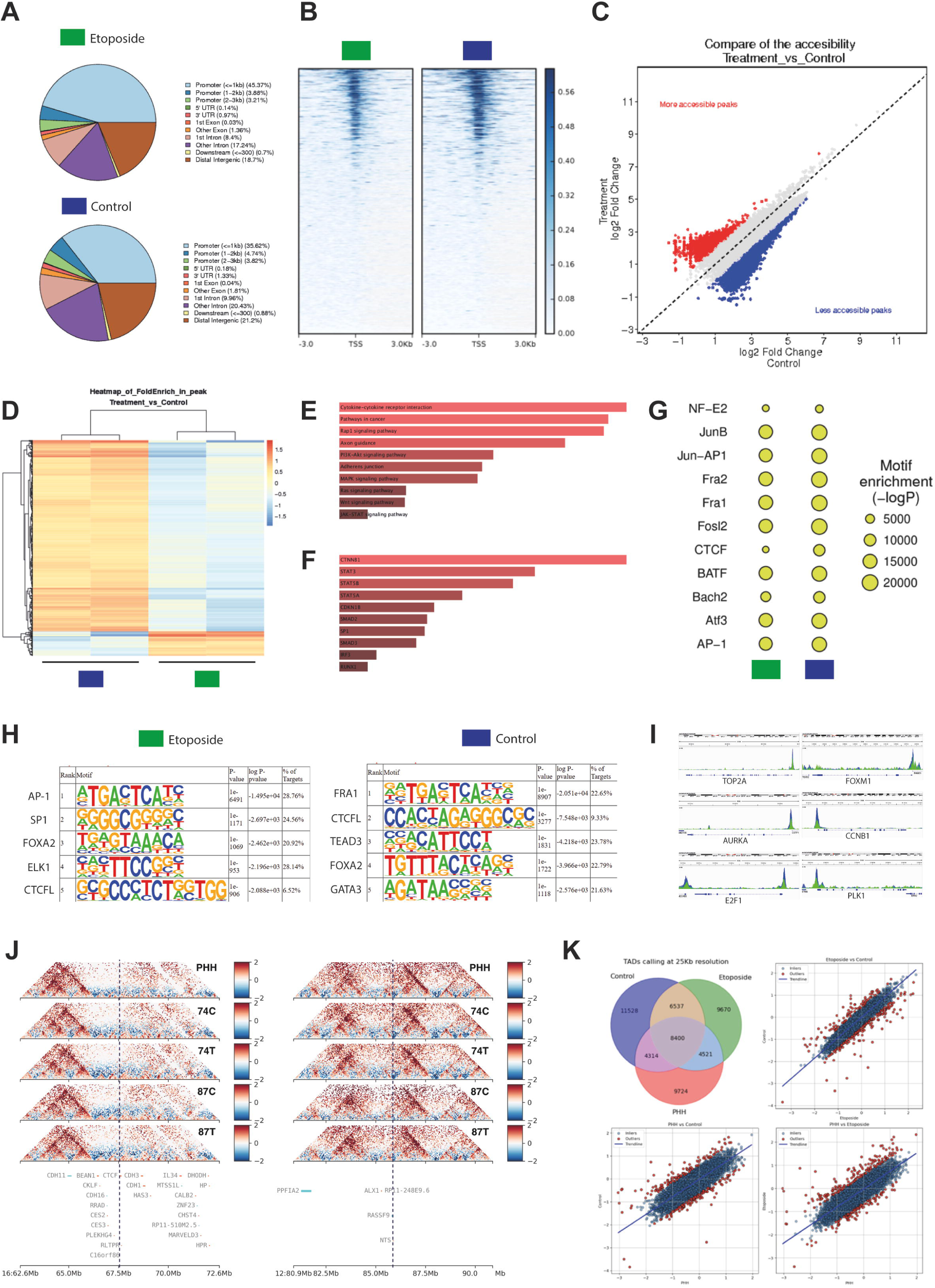
**ATAC-seq and Hi-C analysis of etoposide-treated HLCs**. **A**) Peak distribution in functional gene regions of control and etoposide-treated HLCs. **B**) Distribution of the reads mapped to the TSS. The colour indicates the average density of reads. **C**) Comparison of global chromatin accessibility. Treatment, etoposide-treated HLCs; Control, control HLCs. **D**) Heatmap of z-scored peak fold enrichment between etoposide-treated and control HLCs. The colour represents the average normalized enrichment per chromatin domain. **E**) KEGG analysis of the top 100 enriched peak chromatin domains between etoposide-treated and control HLCs. **F**) Transcription factor protein-protein interaction (PPI) analysis of the top 100 enriched peak chromatin domains between etoposide-treated and control HLCs. **G**) Enrichment of motif and known transcription factor motif. The size of the circle indicates the average degree of enrichment of the motif’s motif and the corresponding transcription factor’s motif. **H**) Motif analysis of the conserved sequence of the peak enrichment position. The motifs are averaged by control or etoposide-treated HLCs and ranked by p-value. **I**) Chromatin accessibility analysis of *TOP2A*, *FOXM1*, *AURKA*, *CCNB1*, *E2F1*, and *PLK1* genes. The peaks represent the average pileup signal for control (blue) or etoposide-treated (green) HLCs. **J**) Chromatin architecture analysis. Hi-C contact maps (observed/expected) at 50 kb resolution for primary human hepatocytes (PHH), control (74C, 87C), and etoposide-treated (74T, 87T) HLCs. Left, *CTCF* genomic region; Right, *NTS* genomic region. **K**) Topologically associating domains (TADs) and insulation scores. Upper left, venn diagram of shared and condition-specific TAD boundaries identified at 25 kb resolution. Upper right and bottom, pairwise comparisons of insulation scores at TAD boundaries between samples. Each point represents an individual TAD boundary; Red dots, outliers with insulation score differences greater than three standard deviations; Blue lines, linear regression trend lines and Pearson correlation coefficient. In every experiment, *n* = 2 biological replicates. Blue, control HLCs; Green, etoposide-treated HLCs. In E and F, the color and length of the bars indicate the combined ranked p-value and odds ratio.

TOP2B interacts with CTCF and influences gene expression through topological structures (26). In HLCs, 10 μM etoposide inhibits TOP2A but not TOP2B, and Hi-C analysis near differentially expressed genes revealed no significant changes in chromatin contacts (Fig. 3J), suggesting that expression changes were independent of genome architecture. TAD and loop analyses demonstrated high similarity across samples, with HLCs and controls exhibiting greater boundary and loop strength concordance (6,537) compared to PHHs (4,314 in control, 4,521 in treated) (Fig. 3K). PHHs displayed stronger loop anchor contacts (p=0.016) (Supplementary Fig. S2B) and more distinct compartmentalization beyond 40 Mb (Supplementary Fig. S2D). These findings indicate that etoposide reduced chromatin accessibility and facilitated transcriptional cell cycle arrest via Wnt signaling, while preserving the PHH 3D landscape regardless of treatment.

### FOXM1 Modulates β-Catenin Signaling and Suppresses Fetal Markers

FOXM1 is responsible for regulating TOP2A expression and interacts with beta-catenin (27). We administered RCM-1, an inhibitor of FOXM1, to HLCs for 5 days (Fig. 4A). RCM-1 is known to reduce proliferation and induce apoptosis (28). The CellTiter-Glo assay indicated that concentrations exceeding 30 µM diminished HLC viability (Fig. 4B). At a concentration of 1 µM RCM-1 (chosen for all subsequent assays), immunofluorescence staining demonstrated a reduction in both FOXM1 and phosphorylated Thr600 FOXM1 (pFOXM1) levels (Fig. 4C). HLCs treated with etoposide exhibited decreased pFOXM1 levels, suggesting impaired nuclear translocation. In HLCs treated with RCM-1, there was a reduction in *TOP2A* expression, but not in *TOP2B* (Fig. 4D). Following treatment with etoposide and RCM-1, the expression of *FOXM1* and *MKI67* decreased. While *ALB* and *HNF4A* levels remained stable, *AFP* expression declined with FOXM1 inhibition. Both treatments led to a decrease in *CTNNB1* expression (Fig. 4E), indicating cell cycle arrest through FOXM1 modulation.

**Fig. 4:**
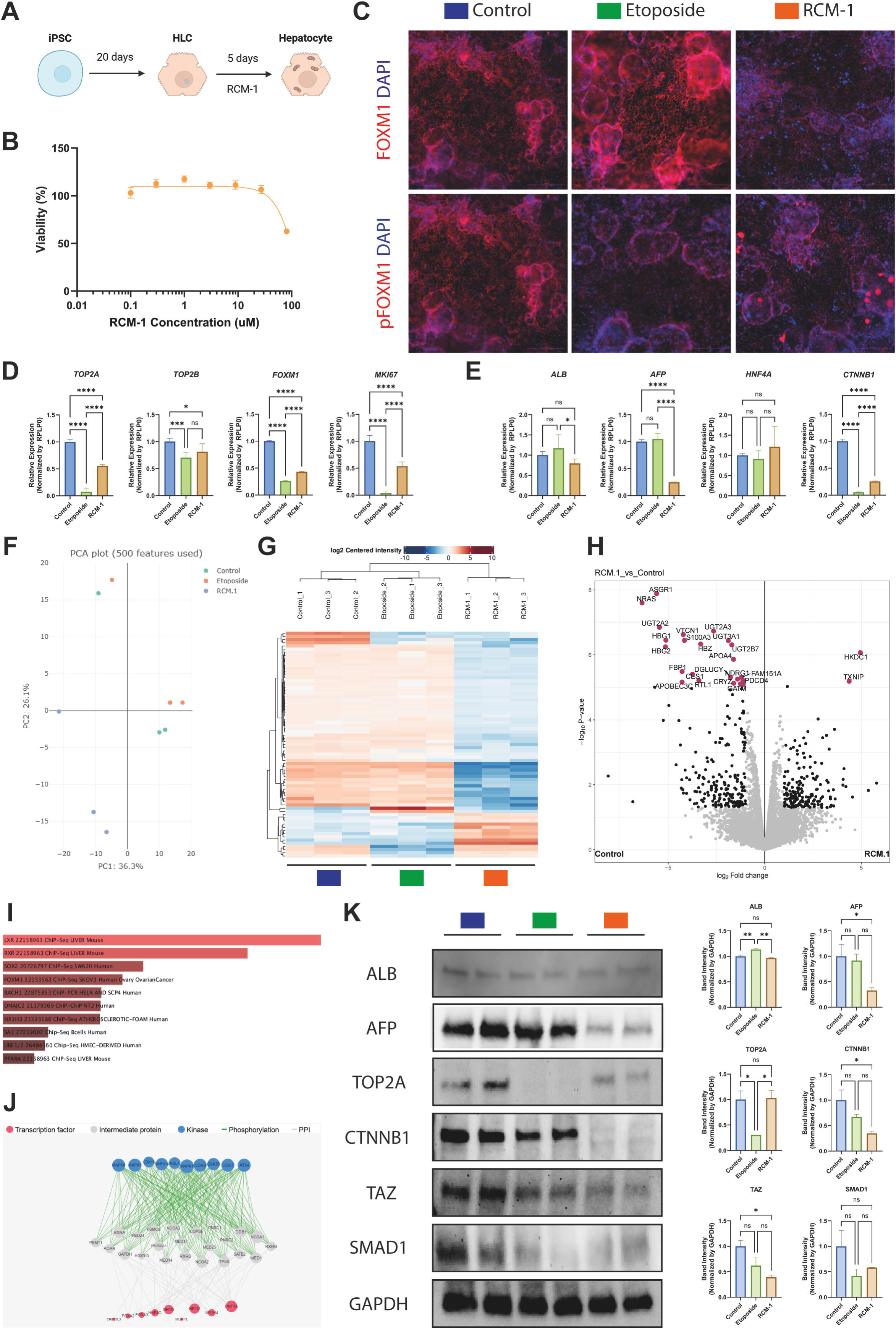
Phosphoproteomic analysis of RCM-1- and etoposide-treated HLCs. **A**) Schematic of RCM-1 treatment of HLCs (Created using BioRender.com). **B**) Cell viability assay of HLCs with different concentrations of RCM-1 to determine toxic doses. **C**) Immunostaining for FOXM1 or phosphorylated Thr600 FOXM1 (pFOXM1) (Red) and DAPI (Blue) (scale bar = 300 μm). **D-E**) RT-qPCR of *TOP2A*, *TOP2B*, *FOXM1*, and *MKI67* (**D**); or *ALB*, *AFP*, *HNF4A*, and *CTNNB1* (**E**) of control, etoposide- and RCM-1-treated HLCs. **F**) Principal component analysis (PCA) on total identified peptides (21,353), adjusted for 500 features. **G**) Heatmap of the top 100 z-scored differentially abundant proteins between all samples. The colour represents the average abundance level of each protein. **H**) Volcano plot of differentially abundant proteins between RCM-1-treated and control HLCs (|log2(FoldChange)| >= 1 & FDR adjusted p-value <= 0.05)**. I**) ChEA analysis of the top 100 differentially abundant proteins between RCM-1-treated and control HLCs. The color and length of the bars indicate the combined ranked p-value and odds ratio. **J**) Expression2Kinases (X2K) analysis of the top 100 differentially abundant proteins between RCM-1-treated and control HLCs. The circle size indicates the number of differentially abundant targets of each transcription factor or kinase. **K**) Representative western blotting gels for ALB, AFP, TOP2A, CTNNB1, YAP/TAZ, SMAD1, and GAPDH. Blue, control HLCs; Green, etoposide-treated HLCs; Orange, RCM-1-treated HLCs. In B, C, D, E, and K, *n* = 2 biological replicates and 3 independent experiments. In F, G, and H, *n* = 3 biological replicates. In B, D, E, and K, data is mean ± SD. In D, E, and K, one-way ANOVA with multiple comparisons and Tukey’s correction (p<0.05).

To further investigate the effects of FOXM1 inhibition, phosphoproteomic analysis revealed that etoposide and RCM-1 formed distinct clusters compared to controls (Fig. 4F). RCM-1 induced a more significant downregulation of proteins compared to etoposide and controls (Fig. 4G). We identified 90 differentially abundant proteins, most of which were down-regulated (Fig. 4H). The top proteins included fetal hemoglobins (HBG1 and HBG2), UGT2B7, and ASGR1, factors expressed in fetal tissues (29, 30, 31). ChEA analysis highlighted enriched LXR and RXR terms (Fig. 4I), crucial for matured hepatocyte lipid metabolism, detoxification, and differentiation (32). The X2K Appyter identified HNF4A/G and FOXA2 as transcription factors; RXRA/B/G and TP53 as intermediate proteins; and GSK3B and ATM as kinases (Fig. 4J). These factors are involved in regulating liver development, proliferation, and differentiation (24, 33, 34), suggesting a role for FOXM1 in liver development pathways.

Next, to validate the qPCR and phosphoproteomic findings, we analyzed the protein abundance using western blotting. FOXM1 inhibition maintained ALB protein levels while reducing AFP levels (Fig. 4K), consistent with gene expression patterns (Fig. 4E). Although RCM-1 reduced TOP2A expression (Fig. 4D), the TOP2A protein levels remained unchanged. β-catenin abundance decreased after TOP2 inhibition, and was depleted by FOXM1 inhibition (Fig. 4K). Wnt signaling enhances YAP/TAZ activity in fetal hepatocytes, whereas the Hippo pathway suppresses it in adult hepatocytes to maintain differentiation. Thus, we evaluated TAZ and SMAD1 protein abundance, identifying a significant reduction of TAZ upon RCM-1 treatment (Fig. 4K). Overall, FOXM1 modulated the β-catenin pathway and suppressed hepatocyte fetal markers, potentially promoting hepatocyte maturation.

### FOXM1 Inhibition Promotes Metabolic Maturation and Polyploidization of HLCs

To assess the long-term effects of TOP2A and FOXM1 inhibition on hepatocyte maturation, HLCs were treated with etoposide and RCM-1 until day 41 of differentiation (D41) (Supplementary Fig. S3A). The lactate dehydrogenase (LDH) release assay revealed no significant cytotoxicity (Supplementary Fig. S3B). In D41 HLCs, *TOP2A*, *FOXM1*, and *MKI67* were significantly downregulated (Supplementary Fig. S3C), indicating cell cycle arrest. *ALB* expression increased with TOP2A inhibition, whereas *AFP* expression was downregulated under both conditions (Supplementary Fig. S3D). *HNF4A* was also significantly downregulated, suggesting the loss of fetal characteristics and induction of adult hepatic function, as the HNF4A P2 isoform decreases postnatally (35). *E2F1* and *E2F8* polyploidization markers that decrease in adult hepatocytes (36) were downregulated by FOXM1 inhibition but not by TOP2A (Supplementary Fig. S3E).

To evaluate functional HLC maturation, we investigated mononucleated and binucleated polyploidization (37). In the human liver, 30% of hepatocytes are polyploid due to cytokinesis failure during early postnatal life (29). Flow cytometry with PI staining revealed that etoposide increased mononucleated polyploidy by 27.6%, while RCM-1 led to a 7.4% increase (Fig. 5A). Then, to assess the tradeoff between hepatocyte proliferation and metabolic maturation, we assessed HLC metabolic maturation upon inducing cell-cycle arrest (38). We found enhanced CYP1A2 and CYP2B6 activities with FOXM1 inhibition (∼2.5×), whereas TOP2 inhibition only increased CYP1A2 (Fig. 5B). The upregulation of CYP1A2 with RCM-1 may be linked to AHR signaling (39, 40), while CYP2B6 upregulation might involve PXR/RXR signaling (41), aligning with our phosphoproteomic findings (Fig. 4I). These pathways are essential for xenobiotic metabolism during terminal differentiation (42). Next, RCM-1 treatment resulted in increased cholesterol secretion (∼1.9×) and decreased triglyceride secretion (∼4.2×) in HLCs (Fig. 5C), indicating enhanced APOB production and effective lipid homeostasis (43, 44). In assessing hepatocyte damage, secreted ALT was higher in controls, while AST levels showed no difference (Fig. 5D). Both ALB and AFP levels decreased during maintenance, with a significant reduction in AFP in treated hepatocytes (Fig. 5E). Western blotting indicated unchanged levels of ALB, CTNNB1, and ERK1/2, but a notable reduction in AFP in RCM-1-treated groups (∼5.6×) (Fig. 5F).

**Fig. 5:**
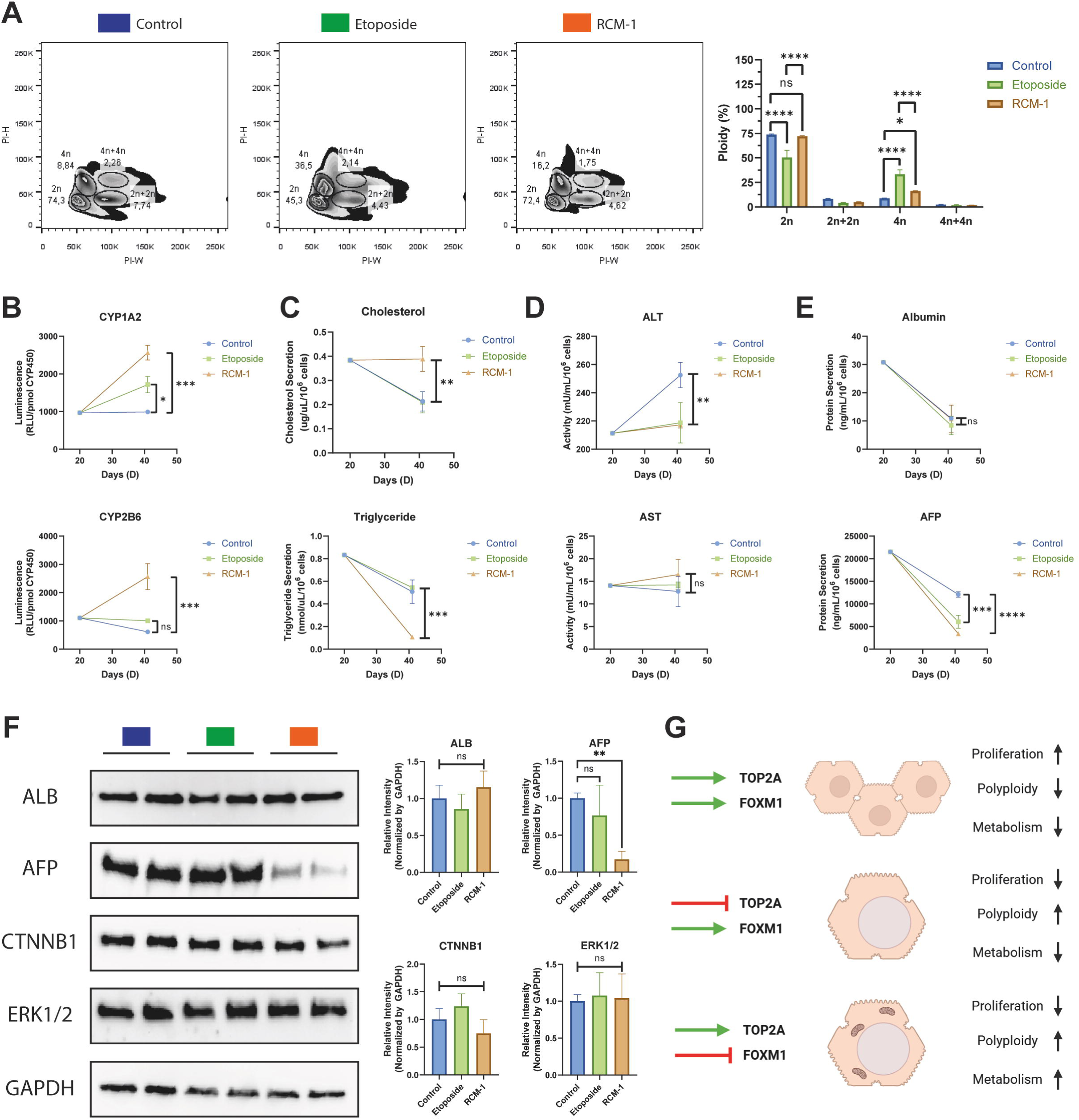
FOXM1 inhibition promotes metabolic maturation and polyploidization of HLCs. **A**) Representative flow cytometry plots of the width versus height of the 535 channel (PI). Mononucleated diploid (2n), mononucleated tetraploid (4n), binucleated tetraploid (2n+2n), and binucleated octoploid (4n+4n) HLCs are shown. **B-E)** ELISA analysis of secreted CYP1A2 and CYP2B6 activity (**B**); secreted cholesterol and triglyceride (**C**); secreted alanine aminotransferase (ALT) and aspartate aminotransferase (AST) activity (**D**); and secreted albumin (ALB) and alpha-fetoprotein (AFP) (**E**). **F**) Representative western blotting gels for ALB, AFP, CTNNB1, ERK1/2, and GAPDH. Blue, control HLCs; Green, etoposide-treated HLCs; Orange, RCM-1-treated HLCs. **G**) Schematic summary for the effects of FOXM1 and TOP2A inhibition in HLCs. Green arrow, activation; Red bar, inhibition (Created using BioRender.com). In every experiment, *n* = 2 biological replicates and 3 independent experiments. In all experiments, data is mean ± SD. In all experiments, one-way ANOVA with multiple comparisons and Tukey’s correction (p<0.05).

In summary, we found that long-term TOP2 and FOXM1 inhibition presented no cytotoxicity and induced mononucleated hepatocyte polyploidization. In addition, only FOXM1 inhibition was associated with HLC metabolic maturation and fetal protein downregulation, potentially indicating consistent in vitro terminal differentiation of hiPSC-derived hepatocytes.

## DISCUSSION

We explored the roles of TOP2A and FOXM1 in the terminal differentiation of HLCs, a critical aspect of liver modeling that current protocols have yet to fully achieve. Inhibiting FOXM1 promoted the terminal differentiation of HLCs by inducing cell cycle arrest, polyploidization, and metabolic maturation, while reducing fetal markers. The transition from fetal to adult stages is governed by the Wnt/β-Catenin Pathway (45, 46). FOXM1 inhibition leads to a decrease in Wnt targets (47) and β-catenin expression, highlighting its significance in hepatocyte maturation. Prolonged FOXM1 inhibition enhanced CYP1A2 and CYP2B6 activities and improves lipid metabolism. miR-122, essential for hepatocyte polyploidization, targets FOXM1 (46), indicating that FOXM1 downregulation may contribute to miR-122-dependent terminal hepatocyte maturation. Extended FOXM1 inhibition resulted in a doubling of mononucleated hepatocyte ploidy, which is linked to the maturation of HLC lipid metabolism. FOXM1 upregulates TOP2A expression by binding to its promoter (48). TOP2A facilitates cell cycle progression during liver development (49), and its downregulation aligns with hepatocyte maturation (50). Long-term inhibition of TOP2A increases HLC polyploidization to levels seen in adult livers and enhances ALB expression, whereas short-term inhibition reduces chromatin accessibility and silences cell cycle genes. Overall, the mutual regulation of TOP2A and FOXM1 may be vital for the late-stage maturation of HLCs.

Although we studied FOXM1 and TOP2A in the terminal differentiation of HLCs, postnatal development creates hepatocyte subpopulations through liver zonation (46). FOXM1 and TOP2A modulation in vitro may produce interzonal-like hepatocytes owing to a lack of metabolic compartmentalization in our system (51). Thus, pericentral, periportal, and interzonal hepatocyte differentiation requires further study. To improve hepatocyte maturation, we proposed targeting FOXM1 and TOP2A using 3D hepatocyte differentiation protocols for a more physiological model, as these enable three-dimensional cell interactions and native extracellular matrix reproduction (52). Our approach was limited to hepatocytes, excluding the interactions between parenchymal and non-parenchymal liver cells. However, iPSCs can produce cholangiocytes, hepatic stellate cells, and Kupffer cells, which are essential for modeling liver disease, and their interactions documented in liver organoid models must be accounted for in development and disease modeling (53). Finally, FOXM1 knockout murine models showed abnormal hepatocyte polyploidization, suggesting that FOXM1 modulation requires fine-tuning during late development (36). Thus, further in vivo studies are needed to validate our findings on FOXM1 and TOP2A modulation of liver development.

Despite our understanding of cell cycle regulation during hepatocyte terminal maturation, we lack clarity regarding how these mechanisms function during embryonic liver development, regeneration, and cancer. Therefore, we hypothesized that FOXM1 was a key candidate for hepatocyte cell cycle regulation. FOXM1 is crucial for hepatocyte cell cycle entry post-hepatectomy, promoting essential proteins such as cyclin A2 and B1 for mitotic progression (54). FOXM1 promotes hepatocyte proliferation and liver mass restoration through its interaction with Wnt/β-catenin signaling (55). Although FOXM1 facilitates regeneration, its overexpression can promote liver cancer (56), which correlates with malignancy and poor prognosis (57). FOXM1 inhibition using small molecules reduces hepatocellular carcinoma growth (58), and experimental models have shown decreased tumorigenic capabilities (59). Thus, investigating FOXM1 modulation may be critical for understanding liver regeneration and carcinogenesis.

In summary, our results indicate that FOXM1 inhibition in HLCs is associated with enhanced hepatocyte terminal differentiation, characterized by cell cycle arrest, polyploidization, and metabolic maturation.

## STATEMENTS

### Author Contributions

K.A.T-S designed and performed the research, analyzed the data, performed bioinformatics analyses, and wrote the paper. L.P., S.K., F.C., G.C., G.C.M., M.G., D.P., V.M.C., and E.G.C. assisted in RT-qPCR, staining, comet, ATAC-seq, phosphoproteomic, and transmission electron microscopy experiments, and revised the manuscript. E.G., M.A., and V.S.F. designed the study and revised the manuscript accordingly. M.Z. designed the study and prepared the manuscript.

