## Supplementary Material for "FOXM1 Inhibition Promotes Polyploidization and Metabolic Maturation in Human iPSC-Derived Hepatocytes by Modulating the Wnt/β-Catenin Pathway"

**Supplementary Data**

1. **Supplementary Tables**

**Supplementary Table 1 - Primer Sequence for RT-qPCR**

| Gene | Forward | Reverse |
|---|---|---|
| *AFP* | CAGGGTGTTTAGAAAACCAGCTAC | TGCAGCAGTCTGAATGTCCG |
| *ALB* | CTCGGCTTATTCCAGGGGTG | AAAGGCAATCAACACCAAGGC |
| *CDKN1A* | TGTCTTGTACCCTTGTGCCTC | CGTTTGGAGTGGTAGAAATCTGTC |
| *CENPF* | AGATGGAGTCCAAGTTGGCG | CGGCCTTGAATAGCATCTTCTG |
| *CTNNB1* | CCTGTTCCCCTGAGGGTATTTG | ACTCCATCAAATCAGCTTGAGTAGC |
| *CYP3A4* | ACCTTGTAAAGAAACACAGATCCC | TCAGGCTCCACTTACGGTG |
| *E2F1* | CAGGAGGTCACTTCTGAGGAGG | GACAACAGCGGTTCTTGCTCC |
| *E2F8* | GGAATTTCGGGCAGCTTCTG | TGAGGCGTTGACACCAAAAAC |
| *FOXA2* | TGCACTCGGCTTCCAGTATG | CATGTTGCTCACGGAGGAGT |
| *FOXM1* | AAACGGGAGACCTGTGATGG | ATCTCTTGCTTGATGCTGCG |
| *GLB1* | TGTGCAGAGTGGGAAATGGG | CTGCCAGGTAATCTGGGTCG |
| *GUSB* | TTCCTATGCCATCGTGTGGG | TGGCGATAGTGATTCGGAGC |
| *HNF4A* | TGGACAAAGACAAGAGGAACC | ATAGCTTGACCTTCGAGTGC |
| *MKI67* | GGATCGTCCCAGTGGAAGAG | GTCTCGTGGGCCACATTTTC |
| *PLK1* | AGTGTCAATGCCTCCAAGCC | TCACAGAGCTGATACCCAAGG |

| | | |
|---|---|---|
| *POU5F1* | AATTTGTTCCTGCAGTGCCC | CACACTCGGACCACATCCTTC |
| *RPLP0* | CATATCCGGGGGAATGTGGG | CAGCAGCTGGCACCTTATTG |
| *TOP2A* | ACCAAGAATCGCCGCAAAAG | TCTCCCCCTTGGATTTCTTGC |
| *TOP2B* | TCTCAGAAGTCAGAAGATGATTCAG | TCTGTTTCAGACCAAATGATGGTG |

**Supplementary Table 2 - Antibodies for Immunostaining**

| Antibody | Catalog Number | Host Species | Assay Dilution |
|---|---|---|---|
| Anti-FOXM1 | 711695 | Rabbit | 1:100 |
| Anti-phospho-FOXM1 (Thr600) | PA5-105625 | Rabbit | 1:100 |
| Goat Anti-Rabbit Alexa 546 | A11010 | Goat | 1:1,000 |

**Supplementary Table 3 - Antibodies for Flow Cytometry**

| Antibody | Catalog Number | Host Species | Assay Dilution |
|---|---|---|---|
| AFP | PA5-21-004 | Rabbit | 1:100 |
| ALB | ab106582 | Chicken | 1:100 |
| FOXA2 | ab60721 | Mouse | 1:100 |
| HNF4A | ab41898 | Mouse | 1:100 |
| OCT3/4 | ab19857 | Rabbit | 1:100 |
| SOX17 | 703063 | Rabbit | 1:100 |
| SSEA4 | ab16287 | Mouse | 1:100 |
| UGT1A1 | ab129729 | Mouse | 1:100 |

| | | | |
|---|---|---|---|
| γH2AX | MA5-33062 | Rabbit | 1:100 |
| Donkey Anti-Mouse Alexa 546 | A10036 | Donkey | 1:1,000 |
| Goat Anti-Mouse Alexa 488 | 11001 | Goat | 1:1,000 |
| Goat Anti-Rabbit Alexa 488 | 11034 | Goat | 1:1,000 |
| Goat Anti-Rabbit Alexa 546 | A11010 | Goat | 1:1,000 |
| Goat Anti-Chicken Alexa 488 | A0001 | Goat | 1:1,000 |

**Supplementary Table 4 - Antibodies for Western Blotting**

| Antibody | Catalog Number | Host Species | Assay Dilution |
|---|---|---|---|
| AFP | ab133617 | Rabbit | 1:1,000 |
| ALB | ab207327 | Rabbit | 1:1,000 |
| CTNNB1 | 9582P | Rabbit | 1:1,000 |
| ERK1/2 | 4695S | Rabbit | 1:1,000 |
| GAPDH | PA1-987 | Rabbit | 1:1,000 |
| SMAD1 | 6944P | Rabbit | 1:1,000 |
| YAP/TAZ | 8418 | Rabbit | 1:1,000 |
| TOP2A | ab219320 | Mouse | 1:1,000 |
| Goat Anti-Rabbit HRP | 7074S | Goat | 1:5,000 |
| Goat Anti-Mouse HRP | 7076S | Goat | 1:5,000 |

2. **Supplementary Methods**

**hiPSC cell maintenance**

All cell culture experiments used three human induced pluripotent stem cells (hiPSCs) from healthy male participants. WTC-11, a characterized control cell line (1), was purchased from the NIGMS Repository. Three other lines (7007, 7405, and 8799) were established at the Human and Stem Cell Research Center. hiPSCs were maintained on 10 µg/cm2 hESC-qualified Matrigel (Corning)-coated plates using Essential 8 Medium (Thermo Fisher) with 100 µg/mL Normocin (InvivoGen). The medium was replaced daily. Cells were split every 3-4 days using Accutase (Gibco), seeded at 3 × 104 cells/cm2 in Essential 8 medium with 5 µM Y-27632 (Sigma-Aldrich) for 24h, and maintained until passage 30 (p30).

**Hepatocyte differentiation**

Hepatocyte differentiation was performed as described previously (2). Single-cell hiPSCs were seeded in Matrigel-coated 6-well plates at 7 × 10^5 cells/well in Essential 8 medium with Y-27632. At 40% confluency on day 2, the medium was changed to RPMI 1640 with 2% B-27 minus Vitamin A 100 ng/mL Activin A 25 ng/mL Wnt3a, and 100 µg/mL normocin. The medium was changed daily for three days and then extended without Activin A two more days. On day 6, the medium was switched to Knockout DMEM with 20% Knockout Serum Replacement, 1% DMSO, 0.5% Glutamax, 1% NEAA, 0.1 mM beta-mercaptoethanol, and 100 µg/mL normocin, and changed daily from days 6 to 8 and on day 10. On day 11, the medium was switched to the Hepatocyte Culture Medium BulletKit with 20 ng/mL oncostatin M and 100 µg/mL normocin, changed every 48h until day 20.

**Transmission electron microscopy**

For transmission electron microscopy (TEM), the cells were dissociated using TrypLE Express (Thermo Fisher Scientific) and pelleted. The Pellets were fixed in 2.5% (v/v) glutaraldehyde in PBS (1x) for 2 h at 4°C, post-fixed in 1% $OsO_4$ for 1 h at 4°C, and stained overnight in 1% aqueous uranyl acetate. The pellets were sequentially dehydrated in 30%, 70%, and 100% ethanol. The samples were then embedded in epoxy resin. Ultrathin sections (70 nm) were obtained using an ultramicrotome, collected on nickel grids, and double stained with uranyl acetate and lead citrate. Micrographs were obtained using a JEOL JEM 1010 electron microscope (80 kV).

**Comet assay**

A comet assay was performed to evaluate genomic DNA damage. The cells were resuspended in agarose, distributed on slides previously coated with the same material, and allowed to solidify for 30 minutes at 4°C. After hardening, the slides were treated with a lysis buffer at 4°C for at least 40 min. After cell membrane lysis and DNA exposure, slides were immersed in an alkaline unwinding solution for 20 min at room temperature. Electrophoresis was conducted in a denaturation buffer at 21 Volts and 0.3 A for 30 min. Staining was performed with SYBR Green (Thermo Fisher Scientific). Images were captured using an Axiovert 200 fluorescence microscope, and 100 nuclei per condition were analyzed using LUCIA Comet Assay software (Laboratory Image). The head/tail DNA ratio (tail momentum) was calculated by using 100 comets on each slide for each sample.

**Hi-C analysis**

The Hi-C library preparation adhered to the Hi-C 2.0 protocol (3). To prepare the Hi-C library, 1–5 × $10^6$ cells were dissociated using trypsin for 15 min, centrifuged at 500 × g for 5 min, washed with PBS, and fixed with 1% paraformaldehyde for 10–12 min. After glycine quenching, cells were pelleted, washed, and stored at − 80°C. Thawed cells were lysed and digested with 300 U of DpnII. The DNA ends were labeled with biotin-14-dCTP using Klenow fragments and ligated with T4 DNA ligase. Crosslinks were reversed using Proteinase K at 65°C, followed by ethanol precipitation. Unligated biotinylated ends were removed using T4 DNA polymerase. NGS libraries were prepared using the KAPA HyperPlus Kit, with biotin-labeled products enriched using Dynabeads MyOne Streptavidin C1, before amplification. Sequencing used 150 bp paired-end reads. Hi-C data were processed using Juicer and Cooltools .hic and .cool files (4). Quality metrics were computed using a modified Juicer script. Differentially expressed gene loci were analyzed using normalized Hi-C maps and were visualized using the FANC library (5). Compartmentalization strength was assessed using cooltools saddle and saddle_strength, with eigenvector decomposition for A/B compartment assignment.

To compare TAD boundary strength between conditions (PHH, control, and Etoposide-treated), we generated scatter plots for each pairwise comparison. Insulation scores were calculated at 25 kb resolution using the cooltools insulation function. The resulting .bed files containing insulation scores were intersected with .bedpe files of TAD boundaries to assign a score to each boundary. For each pairwise comparison, boundaries showing an insulation score difference greater than 3 standard deviations from the main distribution were considered outliers. These outlier regions were subsequently visually validated using Juicebox.

We used two independent sets of chromatin loops in this study. The first set was identified using HiCCUPS (6). Loop length was calculated as the genomic distance between the centroids of loop anchors. For the Venn diagram, loop anchor regions were extended by 10 kb on each side, and pairwise intersections were performed between loops identified in Control, PHH, and Etoposide-treated samples. Loop strength was evaluated using a custom script that calculated the average value of pixels in the balanced submatrix around each loop, derived from matrices normalized with cooltools balance. Additionally, z-scores were computed to evaluate the median loop strength in PHH samples deviated from the distribution observed iPSC-derived.  The second set of loops was identified using the cooltools dots, which detect focal enrichments in contact matrices after estimating expected contact frequencies with cooltools expected-cis. Aggregation of contact signals over loop regions was performed with coolpup.py, followed by visualization using plotpup.py.

The contact probability as a function of genomic distance, P(s), was analyzed to assess chromatin scaling lows. Smoothed and aggregated contact frequency curves were computed using cooltools expected-cis with Gaussian smoothing ($\sigma = 0.1$). Log-log plots of P(s) were generated to visualize contact decay profiles, and first derivatives were computed to highlight changes in the scaling slope across genomic distances (7).

**Proteomics**

In addition to the methods described in the main article, we describe further the methods next. Briefly, proteins were denatured in a Thermomixer (Eppendorf, Germany) at 90 °C and 1200 rom for 5 min and combined with 10 µL of Sera-Mag carboxyl modified magnetic particles (Cytiva, USA) suspension in water a 1 mg/mL After incubating under agitation (900 rpm) for 10 minutes, 200 µL of acetone was added to the mixture and the sample plate was kept at -80°C for 2 hours. The sample plate was transferred to a Microlab Prep (Hamilton, USA) in which all the further steps of protein digestion were achieved. The microparticles complex were magnetically captured using a Magnum EX Universal Magnet Plate (Alpaqua, Beverly, MA, USA), supernatants removed from sample wells and protein extracts were reduced and alkylated with 20 mM TCEP and 15 mM iodoacetamide for 30 min. Microparticles were washed twice with 80% ethanol and the protein extracts were digested overnight at 37°C and 500 rpm with 5 µg of gold mass spec grade trypsin (Promega, Madison, USA) in 100 µL of 100 mM ammonium bicarbonate and 0.01% n-dodecyl-beta-maltoside. The digestion was stopped with the addition of 5 µL of 10% TFA in water, 80% of the trypsin extracts were reserved for phosphopeptides enrichment and remaining volumes were transferred to a Protein LoBind Deepwell plate 96/500 µL and stored at −80 °C until analysis by liquid chromatography-tandem mass spectrometry (LC-MS/MS).

Phosphopeptide elution was achieved in two steps with 2.5% and one with 10% ammonium hydroxide in 50% acetonitrile. Combined extracts were acidified with TFA and concentrated in a SpeedVac vacuum dryer. The chromatograph was equipped with a PepMap100 C18 5 µm, 0.3 × 5 mm trap column and an Aurora Elite 1.7 µm, 75 µm × 15 cm analytical column (Ion Opticks, Australia). Five microliters of tryptic extracts were injected into the LC-MS/MS with 0.1% TFA in water as the transport liquid, at a flow rate of 50 μL/min and trapped for 1 minute. Chromatographic separation was achieved with a binary gradient of 0.1% formic acid in LC/MS-grade water and 0.1% formic acid and 1% DMSO in LC/MS-grade acetonitrile at 0.4 μL/min of flow rate. The content of the organic mobile phase was ramped from 5 to 30% over 29 minutes, then 30 to 50% over 5 minutes, and finally to 90% for 2 minutes. The spectral acquisition was in the data-independent acquisition (DIA) mode achieved by alternating full-scan from m/z 300 to 1200 with a mass resolution of 60,000, followed by 28 MS/MS scans with precursor isolation windows of 25 m/z, with a resolution of 15,000, using standard AGC and maxIT values.

**Immunostaining**

For immunostaining, cells were fixed with 4% paraformaldehyde for 15 min and washed three times with PBS (1x). The samples were permeabilized with 0.01% Triton X-100 in PBS (1x) for 30 min, followed by blocking with 5% bovine serum albumin (BSA) in PBS (1x) for 1 h. The samples were incubated with primary antibodies (listed in Supplementary Table 2) at 4 °C overnight with gentle shaking. The samples were then incubated with secondary antibodies (Supplementary Table 2) at 4 °C for 1 h at room temperature. Nuclei were counterstained 4°Cth DAPI (Sigma) for 1 min at room temperature. Samples were imaged using a Zeiss LSM 800 confocal microscope (Zeiss). Images were analyzed using ImageJ software (v2.0.0).

**Western blotting**

For western blotting, cells were lysed using RIPA Buffer with Protease and Phosphatase Inhibitor Cocktail (Thermo Fisher). Proteins were quantified using a Pierce BCA Protein Assay Kit. Twenty micrograms of protein were denatured using NuPAGE LDS Sample Buffer at 70°C for 10 min. Precision Plus Protein Western C standards were used as size standards. Proteins were loaded onto Novex Tris-Glycine Mini Protein Gels at 120 V for 1 h and then dry-transferred to iBlot 2 PVDF membranes using an iBlot2 Gel Transfer Device. The membranes were blocked with 5% bovine serum albumin (BSA) in TBST for 1 h at room temperature. The membranes were incubated with primary antibodies (Supplementary Table 4) at 4°C overnight using GAPDH as the endogenous control. After washing with TBST, the membranes were incubated with horseradish peroxidase-conjugated secondary antibodies for 1 h. Chemiluminescence was detected using the Amersham ECL Prime Reagent. Protein bands were imaged using the ChemiDoc MP System and analyzed using the ImageLab software.

**Metabolite assays**

Total cholesterol and triglyceride secretion was measured by collecting 1 mL of the supernatant from HLCs cultured in HCM and stored at −80 °C until use. The supernatant was assayed using the HDL and LDL/VLDL Cholesterol Assay Kit (Abcam, ab65390) and Triglyceride Assay Kit (Abcam, ab65336) following the manufacturer's instructions.

**Protein secretion and metabolic activity assays**

Human ALB and AFP secretion were measured by collecting 1 mL of the supernatant from HLCs cultured in HCM and stored at −80 °C until use. The supernatant was assayed using the Human Albumin ELISA Kit (Thermo Fisher, EHALB) and Human Alfa Fetoprotein ELISA Kit (Abcam, ab108838) according to the manufacturer's instructions. Similarly, metabolic activity was measured using the Alanine Transaminase (ALT) Activity Assay (Abcam, ab105134), Aspartate Aminotransferase (AST) Activity Assay (Abcam, ab105135), P450-Glo CYP1A2 Induction/Inhibition Assay (Promega, V8421), and P450-Glo CYP2B6 Assay (Promega, V8321), according to the manufacturer's instructions.

3. **Supplementary Figures Legend**

**Fig. S1: Differentiating iPSCs into HLCs. A**) Schematic for iPSC differentiation into HLCs. iPSC, induced pluripotent stem cell; DE, definitive endoderm; HB, hepatoblast; HLC, hepatocyte-like cell. **B**) Brightfield images of differentiation progression from iPSCs to HLCs (scale bar = 400 μm). **C**) Stepwise flow cytometry characterization of iPSC differentiation into

HLCs for hiPSC markers OCT3/4 and SSEA4; pan-endoderm markers FOXA2 and SOX17; hepatoblast markers AFP and HNF4A; and pan-hepatocyte markers ALB and UGT1A1. Blue, negative isotype-stained control; Red, cell from the respective differentiation stage ($n = 2$, biological replicates and 3 independent experiments). **D**) Stepwise RT-qPCR characterization of iPSC differentiation into HLCs for the iPSC marker *POU5F1*; the endoderm marker *FOXA2*; the hepatoblast markers *AFP* and *HNF4A*; and the hepatocyte markers *ALB* and *CYP3A4* (Data is mean ± SD; ****p<0.001; $n = 2$, biological replicates and 3 independent experiments). In D, one-way ANOVA with multiple comparisons and Tukey's correction.

**Fig. S2: Hi-C analysis of etoposide-treated HLCs. A**) Loop characteristics and overlap across PHH, control, and etoposide-treated samples. Left, distribution of loop lengths (measured as the genomic distance between loop anchors) for each condition. Right, Venn diagram of overlapping chromatin loops across conditions, using a 10 kb tolerance for anchor matching. **B**) Aggregate loop signal and strength. Left, boxplot of loop strength values computed from balanced contact matrices ($z = 2.14$, $p = 0.016$). Right, average pileup plots of chromatin loops. **C**) Contact probability decay curves and their derivatives for aggregated chromosomes. Left, P(s) plots showing the probability of chromatin contacts as a function of genomic separation. Right, first derivative of the log-transformed P(s) curves. **D**) Compartmentalization strength profiles in 100 Kb resolution. **E**) Compartmentalization between PHH and HLCs at different genomic distances. Left, contact maps of chromosome 3 from PHH vs 74T. Right, saddle strength profiles for mid-range interactions (0–20 Mb) and long-range interactions (>40 Mb) at 100 kb resolution. 74C and 87C, vehicle controls; 74C and 87T, etoposide-treated HLCs; PHH, primary human hepatocytes ($n = 2$, biological replicates; data displayed as mean ± SD).

**Fig. S3: Characterization of long-term exposure to RCM-1 and etoposide. A**) Schematic of long-term treatment of HLCs with etoposide or RCM-1. **B**) Cytotoxicity assay (LDH release) of HLCs with RCM-1, etoposide, or 1% DMSO (control) for 41 days. **C-E**) RT-qPCR of *TOP2A*, *TOP2B*, *FOXM1*, and *MKI67* (**C**); *ALB*, *AFP*, *HNF4A*, and *CTNNB1* (**D**); or *E2F1* and *E2F8* (**E**) of control, etoposide- and RCM-1-treated HLCs. In every experiment, $n = 2$ biological replicates and 3 independent experiments. In all experiments, data is mean ± SD. In all experiments, one-way ANOVA with multiple comparisons and Tukey's correction ($p<0.05$).

4. Ma W, Ay F, Lee C, Gulsoy G, Deng X, Cook S, Hesson J, Cavanaugh C, Ware CB, Krumm A, Shendure J, Blau CA, Disteche CM, Noble WS, Duan Z. Using DNase Hi-C techniques to map global and local three-dimensional genome architecture at high

resolution. Methods. 2018 Jun 1;142:59-73. doi: 10.1016/j.ymeth.2018.01.014. Epub 2018 Jan 31. PMID: 29382556; PMCID: PMC5993575.

5. Kruse K, Hug CB, Vaquerizas JM. FAN-C: a feature-rich framework for the analysis and visualisation of chromosome conformation capture data. Genome Biol. 2020 Dec 17;21(1):303. doi: 10.1186/s13059-020-02215-9. PMID: 33334380; PMCID: PMC7745377.

6. Filippova D, Patro R, Duggal G, Kingsford C. Identification of alternative topological domains in chromatin. Algorithms for Molecular Biology [Internet]. 2014 [cited 2025 Apr 30];9:1–11. Available from: https://almob.biomedcentral.com/articles/10.1186/1748-7188-9-14.

7. Durand NC, Shamim MS, Machol I, Rao SSP, Huntley MH, Lander ES, et al. Juicer provides a one-click system for analyzing loop-resolution Hi-C experiments. Cell Syst [Internet]. 2016 [cited 2025 Apr 30];3:95. Available from: https://pmc.ncbi.nlm.nih.gov/articles/PMC5846465/.

### 5. Supplementary Figures

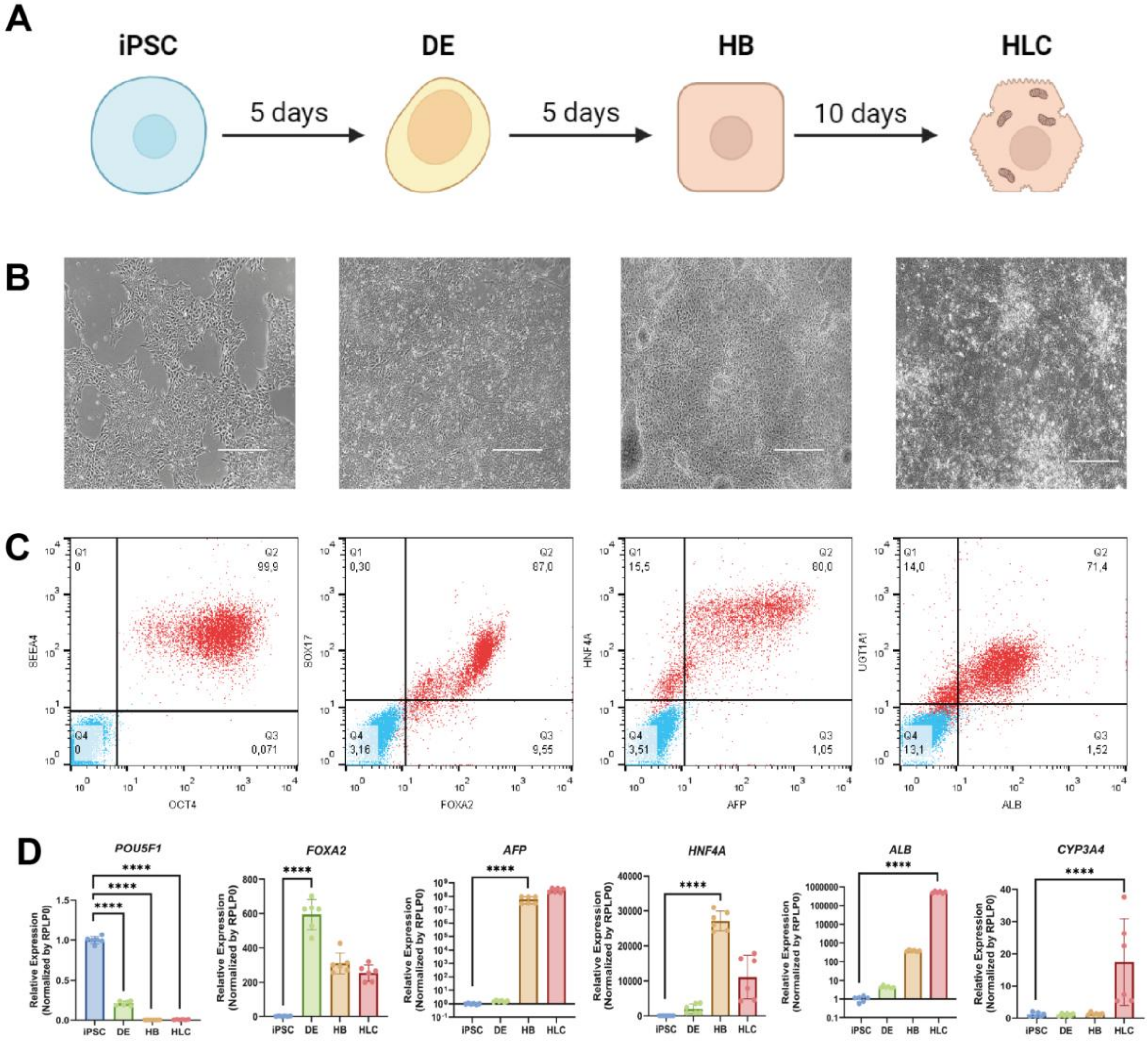
A
iPSC
DE
HB
HLC
5 days
5 days
10 days
B
C
Q1 0
Q2 99,9
Q4 0
Q3 0,071
SEEA4
OCT4
Q1 0,30
Q2 87,0
Q4 3,16
Q3 9,55
SOX17
FOXA2
Q1 15,5
Q2 80,0
Q4 3,51
Q3 1,05
HNF4A
AFP
Q1 14,0
Q2 71,4
Q4 13,1
Q3 1,52
UGT1A1
ALB
D
POU5F1
FOXA2
AFP
HNF4A
ALB
CYP3A4
Relative Expression (Normalized by RPLP0)
iPSC
DE
HB
HLC
****

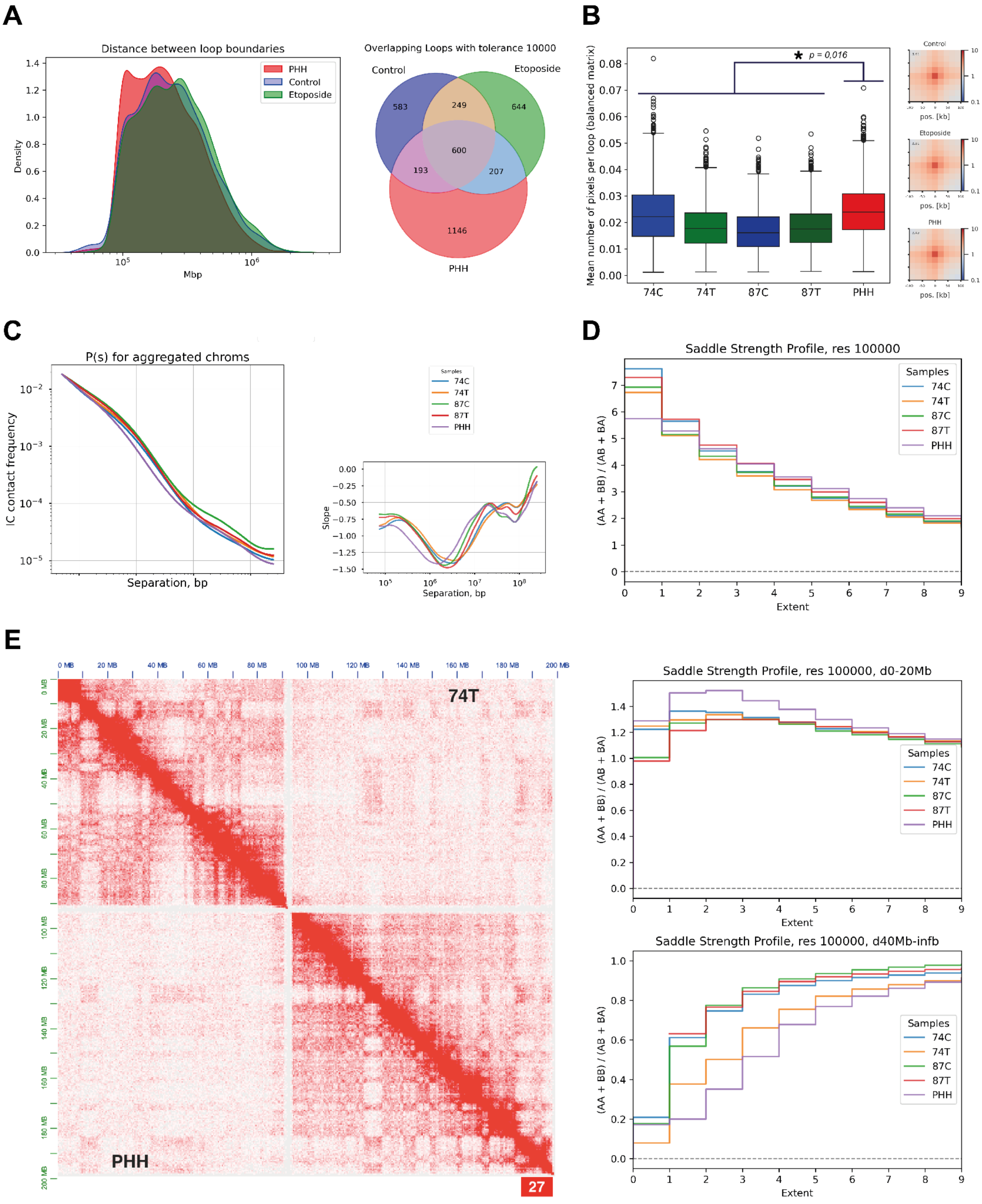
A
Distance between loop boundaries
PHH
Control
Etoposide
Density
Mbp
Overlapping Loops with tolerance 10000
Control
Etoposide
PHH
583
249
644
600
193
207
1146
B
Mean number of pixels per loop (balanced matrix)
p = 0,016
74C
74T
87C
87T
PHH
Control
Etoposide
PHH
pos. [kb]
C
P(s) for aggregated chroms
IC contact frequency
Separation, bp
Samples
74C
74T
87C
87T
PHH
Slope
D
Saddle Strength Profile, res 100000
(AA + BB) / (AB + BA)
Extent
E
74T
PHH
27
Saddle Strength Profile, res 100000, d0-20Mb
Saddle Strength Profile, res 100000, d40Mb-infb

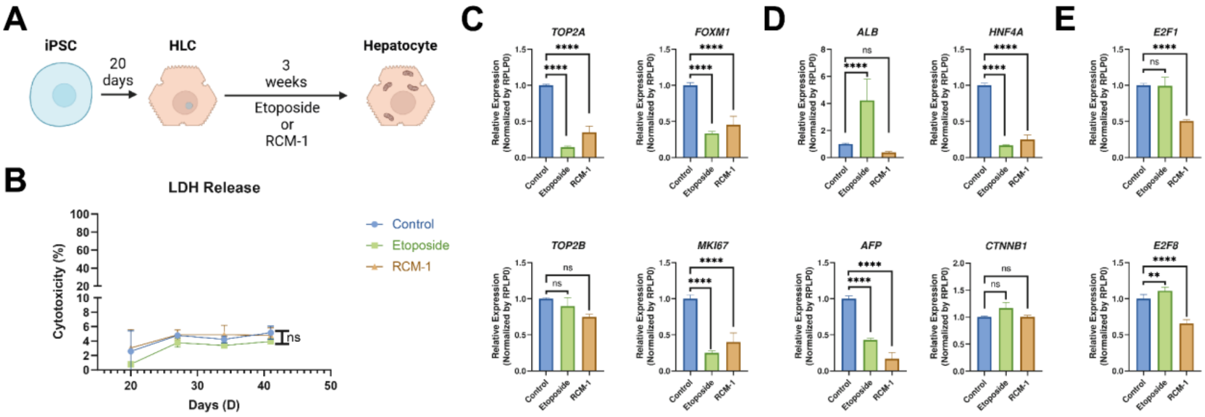

A
iPSC
20 days
HLC
3 weeks
Etoposide or RCM-1
Hepatocyte
B
LDH Release
Cytotoxicity (%)
Days (D)
Control
Etoposide
RCM-1
ns
C
TOP2A
FOXM1
TOP2B
MKI67
D
ALB
HNF4A
AFP
CTNNB1
E
E2F1
E2F8
Relative Expression (Normalized by RPLP0)
Control
Etoposide
RCM-1
